# A Molecular Link between Cell Wall Modification and Stringent Response in a Gram-positive Bacteria

**DOI:** 10.1101/622340

**Authors:** Surya D. Aggarwal, Saigopalakrishna S. Yerneni, Ana Rita Narciso, Sergio R. Filipe, N. Luisa Hiller

## Abstract

To ensure survival during colonization of the human host, bacteria must successfully respond to unfavorable and fluctuating conditions. This study explores the fundamental phenomenon of stress response in a gram-positive bacterium, where we investigate the ability of a cell wall modification enzyme to modulate intracellular stress and prevent the triggering of the stringent response pathway. The *Streptococcus pneumoniae* cell wall modification proteins MurM and MurN are tRNA-dependent amino acid ligases, which lead to the production of branched muropeptides by generating peptide crossbridges. In addition, MurM has been proposed to contribute to translation quality control by preferentially deacylating mischarged tRNAs mischarged with amino acids that make up the peptidoglycan. Here, we demonstrate that the *murMN* operon promotes optimal growth under stressed conditions. Specifically, when grown in mildly acidic conditions, a *murMN* deletion mutant displays early entry into stationary phase and dramatically increased lysis. Surprisingly, these defects are rescued by inhibition of the stringent response pathway or by enhancement of the cell’s ability to deacylate mischarged tRNA molecules. The increase in lysis results from the activity of LytA, and experiments in macrophages reveal that *murMN* regulates phagocytosis in a LytA-dependent manner. These results suggest that under certain stresses, these bacterial cells lacking MurMN likely accumulate mischarged tRNA molecules, activate the stringent response pathway, and enter prematurely into stationary phase. Moreover, by virtue of its ability to deacylate mischarged tRNAs while building peptidoglycan crossbridges, MurM can calibrate the stress response with consequences to host-pathogen interactions. Thus, MurM is positioned at the interface of cell wall modification, translation quality control and stringent response. These findings expand our understanding of the functions of the bacterial cell wall: cell wall modifications that impart structural rigidity to the cell are interlinked to the cell’s ability to signal intracellularly and mount a response to environmental stresses.

**SIGNIFICANCE:** During infection, microbes must survive the hostile environmental conditions of the human host. When exposed to stresses, bacteria activate an intracellular response, known as stringent response pathway, to ensure their survival. This study connects two fundamental pathways important for cellular growth in a gram-positive bacterium; it demonstrates that enzymes responsible for cell wall modification are connected to the stringent response pathway via their ability to ameliorate errors in protein translation. Our study was performed on *Streptococcus pneumoniae* where the cell wall modification enzyme, MurM, is a known determinant of penicillin resistance. We now demonstrate the importance of MurM in translation quality control and establish that it serves as a gatekeeper of the stringent response pathway.

## INTRODUCTION

Gram-positive bacteria have evolved a thick and sophisticated cell wall that ensures bacterial structural integrity and is critical for cellular viability. This dynamic structure serves as a scaffold for surface anchored proteins, wall teichoic acids, lipoteichoic acids, lipoproteins, and capsular polysaccharides (1, 2). It is also a major target of immune defenses and antibiotics (3, 4). Opportunistic pathogens, such as *Streptococcus pneumoniae* (pneumococcus), encounter hostile environments within the human host, where the cell wall serves as a barrier and an interface between the bacteria and its host.

The pneumococcal cell wall consists of glycan chains that are cross-linked via peptide bridges (5). These glycan chains consist of alternating subunits of *N*-acetylglucosamine (NAG) and *N*-acetylmuramic acid (NAM) residues. Each NAM subunit is attached to a peptide side chain, which is crosslinked to stem peptide from another glycan chain. In pneumococcus, the link can be direct or through a short diamino acid peptide bridge that is assembled by MurM and MurN, two tRNA-dependent amino acid ligases. These proteins add the diamino acid peptide to the lipid II peptidoglycan precursor in the inner side of the bacterial membrane before this complex is flipped to the outer side, where the peptidoglycan building block containing this diamino acid peptide is incorporated into the cell wall by penicillin binding proteins (6). The peptide bridge originates from the third residue (L-Lys) on one pentapeptide chain and is ultimately connected to the fourth residue (D-Ala) of the stem peptide in a neighboring glycan chain (6, 7). MurM contributes either an alanine or a serine to the first position of the bridge, while MurN contributes an alanine to the second position. Thus the pneumococcal peptidoglycan peptide bridge consists of either L-Ala-L-Ala or L-Ser-L-Ala. Allelic variants of *murM* alone lead to diversity in the nature of stem peptide branching in the cell wall peptidoglycan (8–10). Finally, *murMN* are required for penicillin resistance, as inactivation of this operon leads to a complete loss of penicillin resistance (6, 11).

As tRNA-dependent amino acid ligases, MurMN acquire the alanines and serines they utilize for the peptidoglycan bridge from aminoacyl-tRNAs. The employment of charged tRNAs as substrates for MurMN is an evolutionary design that juxtaposes the fundamental cellular functions of peptidoglycan biosynthesis and protein translation. Previous work has demonstrated that MurM can deacylate mischarged tRNAs (12). Moreover, *in vitro*, it displays a substantial preference for deacylating mischarged tRNAs over correctly charged tRNAs (13). The preference for amino acids from mischarged tRNAs strongly suggests that MurM decreases the pool of mischarged tRNAs present in the cell, thereby, contributing to the process of translation quality control (12).

Under stress, many bacterial cellular processes including translation, are regulated by the stringent response pathway. Deprivation of amino acids or carbon sources, elevated temperature, and acidic conditions can trigger the stringent response and rewire the cellular circuitry to adapt to challenging conditions and ensure bacterial survival (14–17). In addition, the stringent response plays a role in regulation of bacterial virulence and susceptibility to antimicrobials (15, 18–20). The stringent response pathway is characterized by the accumulation of alarmones, specifically guanosine tetra- (ppGpp) and pentaphosphate (pppGpp). In the majority of gram-negative bacteria, these are produced by RelA. In gram-positive bacteria, these are produced by a bifunctional RSH protein (RelA/SpoT homolog), with both synthetase and hydrolase activity (16, 21). In gram-negative bacteria, binding of deacylated-tRNA to ribosomes activates alarmone production (22–25), however, the molecular mechanisms underlying activation of the stringent response pathway in gram-positive bacteria remain elusive.

In this study, we show that MurMN is a molecular link between the processes of cell wall modification and translation quality control. Our findings indicate that the absence of these proteins sensitizes pneumococcal cells to acidic stress: when a *murMN* deletion mutant strain is grown in mildly acidic conditions, growth defects result from the likely accumulation of mischarged tRNAs and the activation of the stringent response pathway. Further, our data suggest that activation of the stringent response stimulates LytA activity and subsequently influences pneumococcal cells’ ability to be phagocytosed. These findings provide insight into cell wall function by suggesting that the cell wall modification enzymes can buffer intracellular stress and modulate activation of the stringent response pathway.

## RESULTS

### The *murMN* operon protects cells against acid-induced growth defects

When growing a *ΔmurMN* strain in planktonic culture, we observed a pronounced growth defect in rich media in mildly acidic conditions (pH 6.6) (**Fig. 1A**). The mutant exhibited increased susceptibility to stationary-phase induced autolysis and a decrease in the maximum optical density reached after growth (max OD), consistent with an early onset of stationary phase. This growth defect was rescued in a *ΔmurMN* complemented strain (*ΔmurMN*:*murMN*) (**Fig. 1B and 1C**). Although minor differences in growth of wild-type and *ΔmurMN* strains were observed in rich media at normal conditions (pH 7.4), these differences were much more pronounced in mildly acidic conditions.

**Fig. 1.**
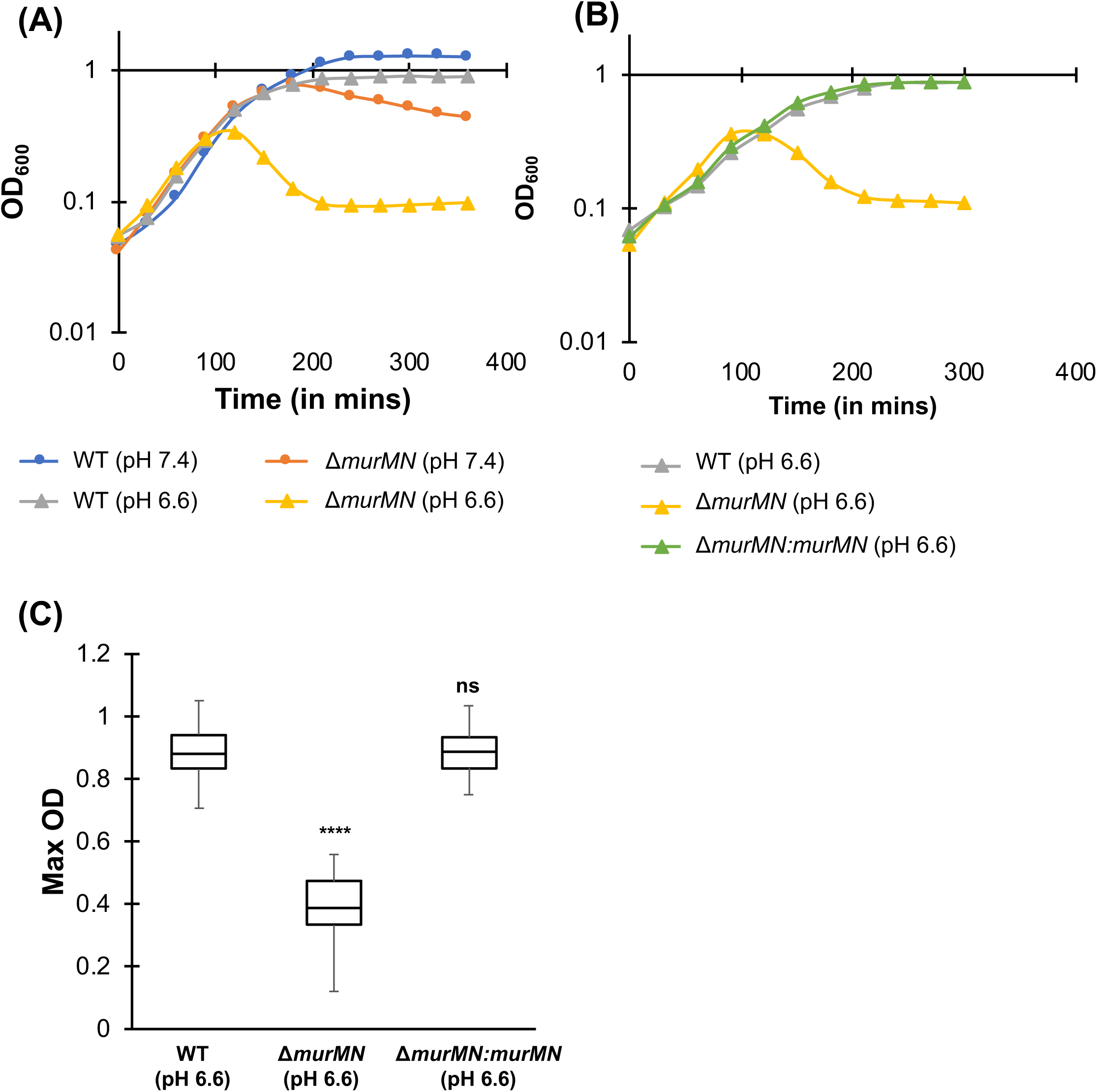
Growth defects observed in the absence of *murMN*. **(A)** Representative growth curves for wild-type and *ΔmurMN* strains grown in normal (●) and acidic conditions (▲). Growth curves were started at an OD_600_ of 0.05 *(at least n=3)*. **(B)** Representative growth curves for wild-type, *ΔmurMN* and *ΔmurMN:murMN* strains grown in acidic conditions (▲). Growth curves were started at an OD_600_ of 0.05 *(at least n=3)*. **(C)** Box-and-whisker plot depicting median and range of maximum growth of wild-type, *ΔmurMN* and *ΔmurMN:murMN* strains grown in acidic conditions. ‘ns’ denotes non-significant comparison and **** *p*<0.0001 relative to wild-type strain by ANOVA followed by Tukey’s post-test.

MurMN contribute to cell wall branching by adding diamino acid peptide bridges to the peptidoglycan stem peptide that lead to production of branched muropeptides (6). The loss of branching alters pneumococcal cell wall composition and increases bacterial sensitivity to lysis by cell wall inhibitors (26). Thus, one hypothesis for the difference in sensitivity of wild-type and *ΔmurMN* strains to acidic stress is that wild-type cells incur changes in the cell wall in response to pH, and that these chances are absent in a *ΔmurMN* strain. To test this hypothesis, we employed high-performance liquid chromatography (HPLC) to analyze the peptidoglycan composition of pneumococci grown in rich media at normal (pH 7.4) and mildly acidic (pH 6.6) conditions for both wild-type and *ΔmurMN* strains. We did not observe any differences in the cell wall composition of the wild-type cells between normal and acidic conditions (**Fig. 2Ai, B**). Similarly, there was also no difference in the cell wall composition of *ΔmurMN* cells between normal and acidic conditions (**Fig. 2Aii, B**). As expected, we observed that deletion of *murMN* led to the loss of branched peptides from the peptidoglycan compared to wild-type cells. These findings suggest that the growth differences observed in the *ΔmurMN* strain in acidic conditions cannot be attributed to pH-dependent alterations in peptidoglycan composition.

### Disruption of the translation quality control function of MurM leads to pneumococcal growth defects

Recent *in vitro* experiments have shown that MurM can deacylate mischarged tRNAs, and that it may play a significant role in maintaining low levels of mischarged tRNAs in pneumococcal cells (12). Given that the growth defect of the *ΔmurMN* strain could not be attributed to alterations in the cell wall composition in acidic conditions (**Fig. 2**), we hypothesized that the loss of the translation quality control role of MurM was responsible for the observed phenotypes.

**Fig. 2.**
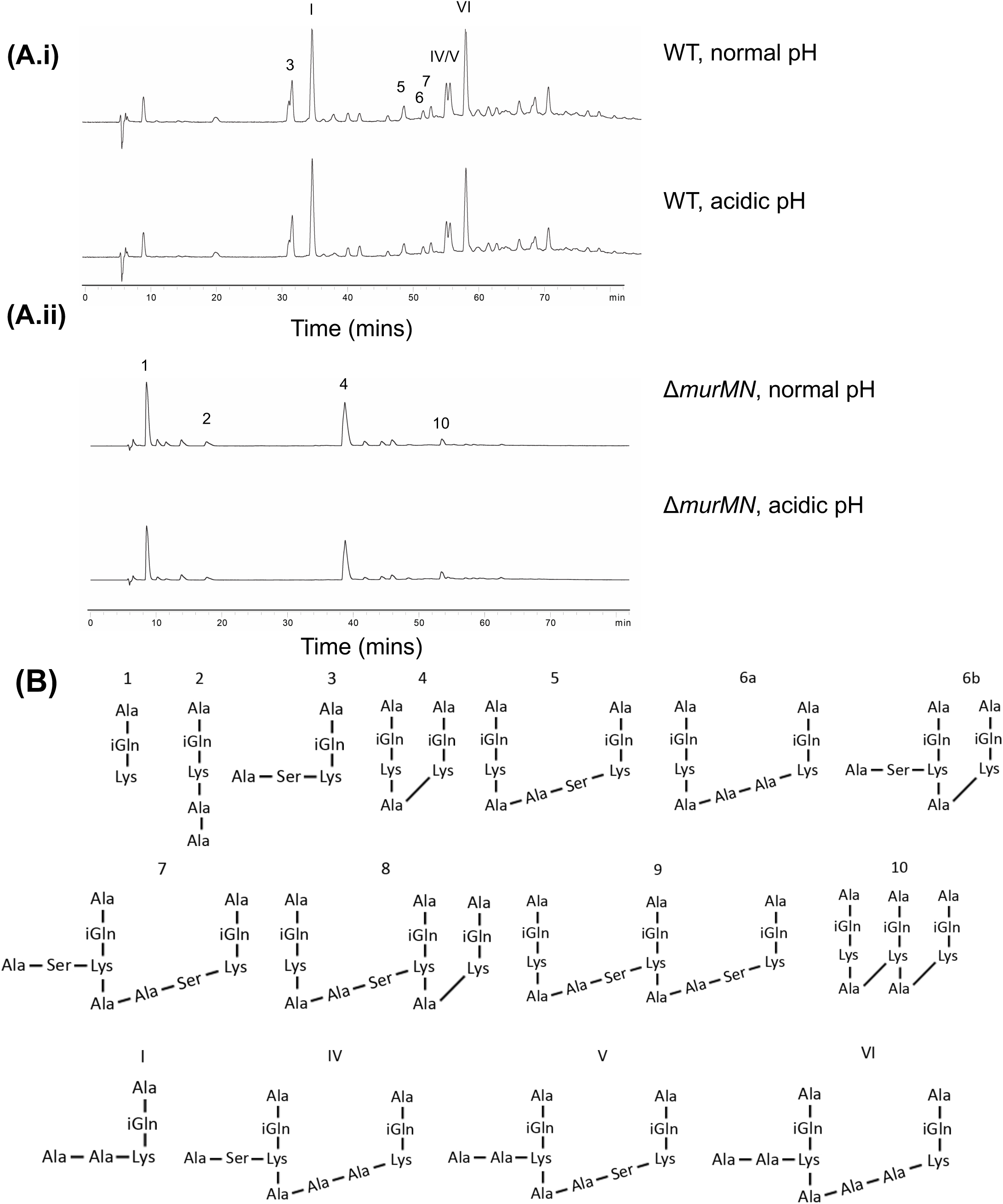
Analysis of peptidoglycan structure of cells grown in normal and acidic conditions. HPLC profiles of the stem peptide compositions of peptidoglycan from **(A.i)** wild-type and **(A.ii)** *ΔmurMN* strains grown in normal and acidic pH. **(B)** Structures of the cell wall stem peptides that comprise pneumococcal peptidoglycans.

In addition to obtaining alanine or serine from correctly aminoacylated-tRNAs (Ala-tRNA^Ala^ or Ser-tRNA^Ser^ respectively), MurM can also acquire these amino acids from other mischarged tRNA species (12, 27). In fact, alanyl-tRNA synthetase (AlaRS), the enzyme responsible for charging tRNA^Ala^ to produce Ala-tRNA^Ala^ is error prone, such that tRNA^Ala^ is often misactivated with serine (Ser-tRNA^Ala^) or glycine (Gly-tRNA^Ala^) (28) (**Fig. 3B**). Accumulation of Ser-tRNA^Ala^ is toxic to cells such that many organisms, across all the three kingdoms of life, encode a free-standing homolog to the editing domain of AlaRS protein, termed AlaXp. These AlaXp prevent toxicity by deacylating mischarged tRNA^Ala^ molecules in different organisms (29–33). Interestingly, analysis of the *S. pneumoniae* available genomes, suggests that pneumococcus does not encode an AlaXp protein; instead, once the tRNA molecules are charged, the responsibility of ensuring translational fidelity has been proposed to rest with MurM by virtue of its deacylation function (12, 34). Importantly, the catalytic efficiency of MurM in using mischarged Ser-tRNA^Ala^ as a substrate for generating crosslinks is dramatically higher relative than its ability to use correctly acylated Ala-tRNA^Ala^ or Ser-tRNA^Ser^ (13). The high affinity of MurM for misacylated-tRNA^Ala^ combined with the absence of a gene encoding AlaXp in the pneumococcal genome, emphasize the importance of MurM in translation quality control.

**Fig. 3.**
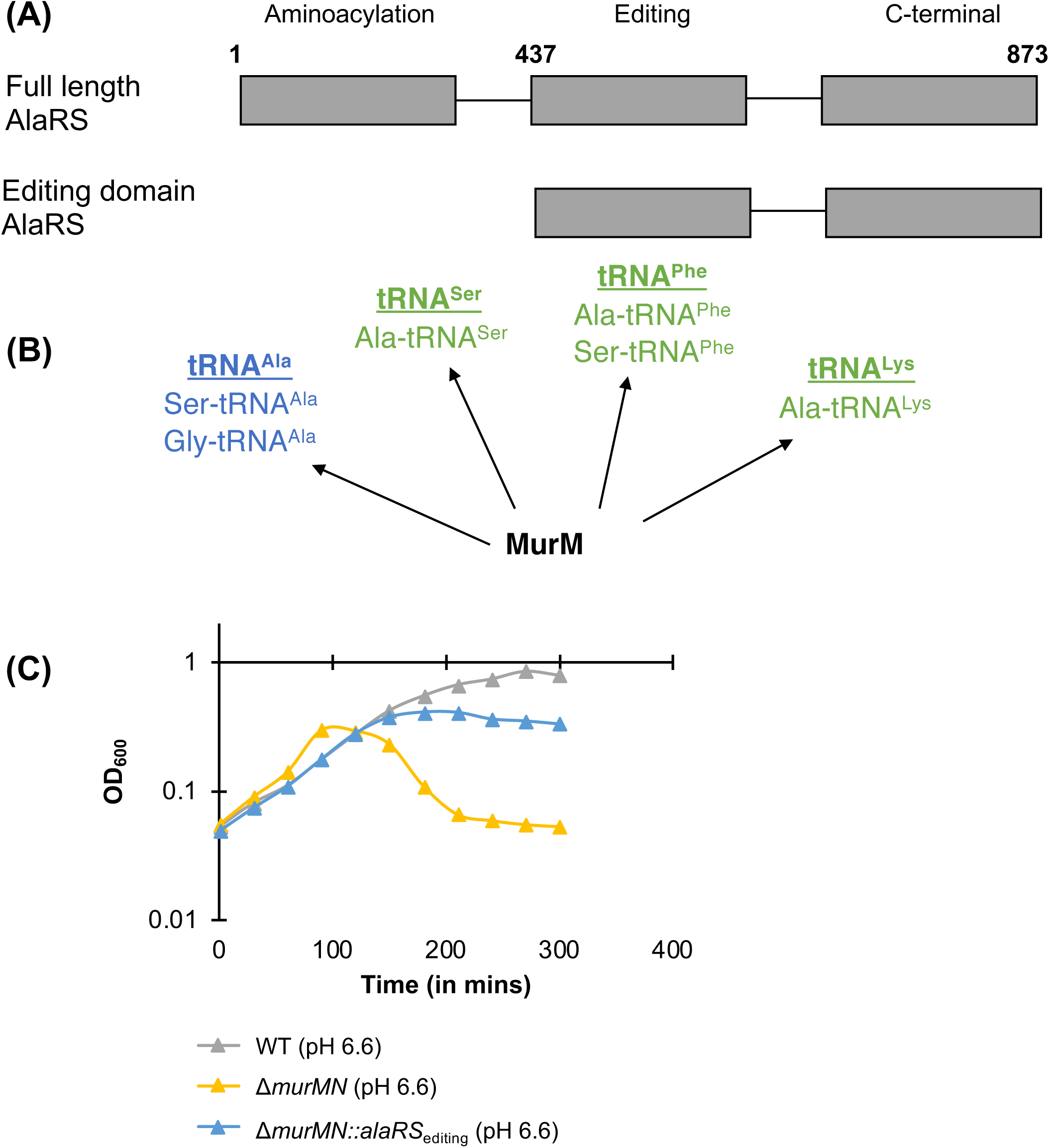
Analysis of translation quality control function of MurM. **(A)** Schematic representation of the domain architecture of full-length and ectopically expressed editing domains of AlaRS. **(B)** Different tRNA moieties that can be deacylated by MurM (underlined). *In blue*: Mischarged tRNA moieties that can be edited by AlaRS_editing_ protein. *In green:* All other mischarged tRNA moieties that can be edited by MurM but not AlaRS_editing._ Schematic based on findings from previous work (12, 27). **(C)** Representative growth curve for wild-type, *ΔmurMN* and *ΔmurMN::alaRS*_editing_ strains grown in acidic conditions (▲). Growth curves were started at an OD_600_ of 0.05 *(at least n=3*).

We reasoned that expression of a tRNA-editing protein in a *ΔmurMN* background could allow us to distinguish between the cell-wall crosslinking and the translation quality control roles of MurM, and thus establish which of these roles is responsible for the growth defect of the *ΔmurMN* strain. To this end, we overexpressed the editing domain of AlaRS protein. We determined the editing domain of the protein encoded by pneumococcal *alaRS* (*spr_1240*) using an amino acid alignment with AlaRS proteins from *E. coli* (WP_096112793.1) and *B. subtilis* (WP_007408211.1), where the protein domains have been well characterized (30, 31, 35). The pneumococcal editing domain was identified as residues 437-873 of the pneumococcal AlaRS, and the peptide encoded by this region was used for functional complementation (AlaRS_editing_) (**Fig. 3A**). Ectopic expression of AlaRS_editing_ is predicted to deacylate mischarged alanine-specific tRNA species (tRNA^Ala^), the most abundant of which are Ser-tRNA^Ala^ and Gly-tRNA^Ala^ (12, 28) (blue in **Fig. 3B**). Since AlaRS_editing_ only deacylates mischarged tRNA^Ala^ molecules, its expression will not diminish the correction of other mischarged tRNA species (tRNA^Ser^, tRNA^Phe^, tRNA^Lys^), which are also thought to be employed by MurM to acquire non-cognate alanine or serine (12, 13, 27) (green in **Fig. 3B**).

The expression of AlaRS_editing_ in the *ΔmurMN* strain partially rescued the growth defect displayed by the *ΔmurMN* strain in mildly acidic conditions (**Fig. 3C**). Specifically, the *ΔmurMN::alaRS*_editing_ strain resembled the wild type in that it did not exhibit the dramatically increased stationary-phase induced lysis associated with the *ΔmurMN* and its maxOD was intermediate between the *ΔmurMN* and wild-type strains. These finding illustrate the importance of the deacylating function of MurM *in vivo* and imply that MurM ensures normal growth in mildly acidic conditions by its ability to deacylate mischarged tRNAs, which would be toxic for pneumococcal cells.

### MurMN regulate activation of the stringent response pathway

The stringent response is a well conserved pathway utilized by bacteria to ensure survival under stressed conditions. Mediated by the production of alarmones, the activation of this pathway rewires the cellular circuitry and coordinates cellular entry into stationary phase (36). We hypothesized that the differences in growth of the *ΔmurMN* strain, relative to the wild-type strain, are a consequence of activation of the stringent response pathway.

If activation of the stringent response pathway contributes to the early onset of stationary phase in the *ΔmurMN* strain, this growth phenotype should be decreased or abrogated in a *ΔmurMN* where the stringent response pathway cannot be activated. This pathway is activated by accumulation of the intracellular alarmones, ppGpp or pppGpp, which are hyperphosphorylated forms of GDP or GTP respectively, synthesized by the addition of a pyrophosphate molecule obtained from ATP (**Fig. 4A**). In pneumococcus, the primary source of alarmone production is RSH (<u>R</u>elA/<u>S</u>poT <u>H</u>omolog), a bifunctional synthetase and hydrolase of (p)ppGpp (37). Thus, we deleted *rsh* (*spr_1487*) to generate strains defective in the ability to activate the stringent response pathway. Thin-layer chromatography experiments were used to confirm that the *Δrsh* strain did not produce measurable levels of alarmone (**Fig. 4B**).

**Fig. 4.**
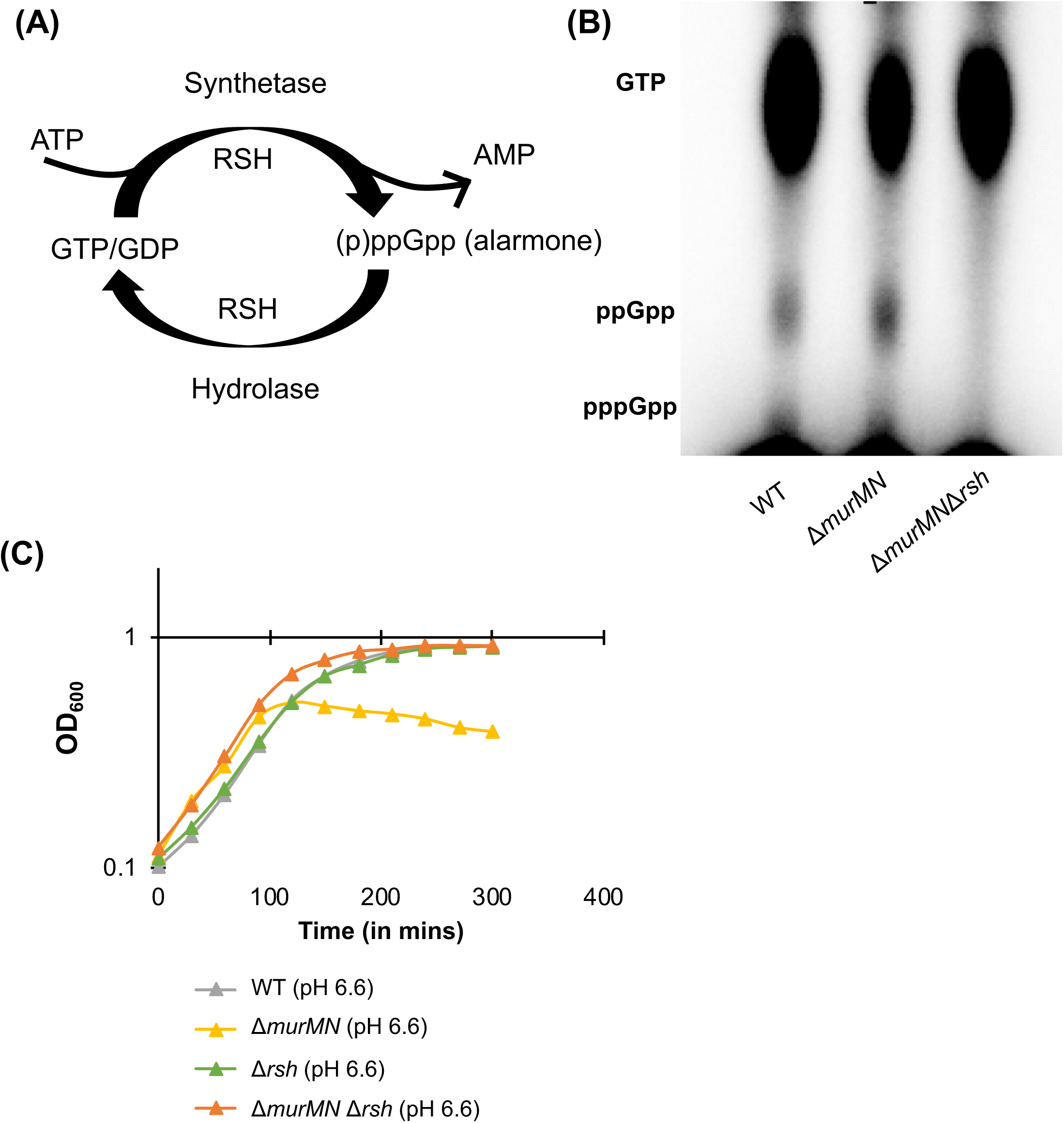
MurMN regulates stringent response pathway. **(A)** Schematic of (p)ppGpp production and hydrolysis. **(B)** Thin-liquid chromatography showing ^32^P being incorporated as GTP or alarmones (ppGpp or pppGpp) in wild-type, *ΔmurMN* and *ΔmurMNΔrsh* strains grown in acidic conditions. Ratio of alarmone to GTP in *ΔmurMN* is 3.7-fold times that in the wild-type. **(C)** Representative growth curve for wild-type, *ΔmurMN*, *Δrsh* and *ΔmurMNΔrsh* strains grown in acidic conditions (▲). To circumvent the long lag phase of strains from *Δrsh* background, strains were grown to log phase (~OD_600_ 0.2), collected by centrifugation and growth curves started at an OD_600_ of 0.1 *(at least n=3)*.

We compared growth among wild-type, *ΔmurMN,* and*ΔmurMNΔrsh* strains (**Fig. 4C**). The *ΔmurMNΔrsh* strain did not exhibit the growth defects observed for the *ΔmurMN* strain in mildly acidic conditions. Thus, inhibition of the stringent response abrogates the growth defects in a *ΔmurMN* background. To measure the relative alarmone levels in the *ΔmurMN* strain relative to the wild-type strain in mildly acidic conditions, we employed thin-layer chromatography (TLC). The *ΔmurMN* strain displays increased alarmone levels compared to the wild-type strain, as observed by the higher levels of alarmone relative to GTP, consistent with growth defects in the absence of *murMN* (**Fig. 4B**). We conclude that acidic conditions promote entry into stringent response, and that MurMN buffer the activation of stringent response and consequently entry into stationary phase.

It is well established that *murMN* is required for penicillin resistance of pneumococci (6). We investigated whether the fitness of *ΔmurMN* strain under penicillin pressure can be enhanced by inhibition of the stringent response pathway. To this end, we determined the percentage of viable pneumococcal cells when grown at a concentration of penicillin, that is approximately equal to the minimum inhibitory concentration of this antibiotic for the *ΔmurMN* strain (0.025 µg/ml). At this concentration, the *ΔmurMNΔrsh* and the *ΔmurMN::alaRS*_editing_ strains displayed a marked increase in survival relative to the *ΔmurMN* background (**Table 1**). We conclude that the role of MurM in preventing the activation of the stringent response, due to correction of mischarged tRNAs, contributes to increase in fitness under penicillin pressure.

**Table 1:**
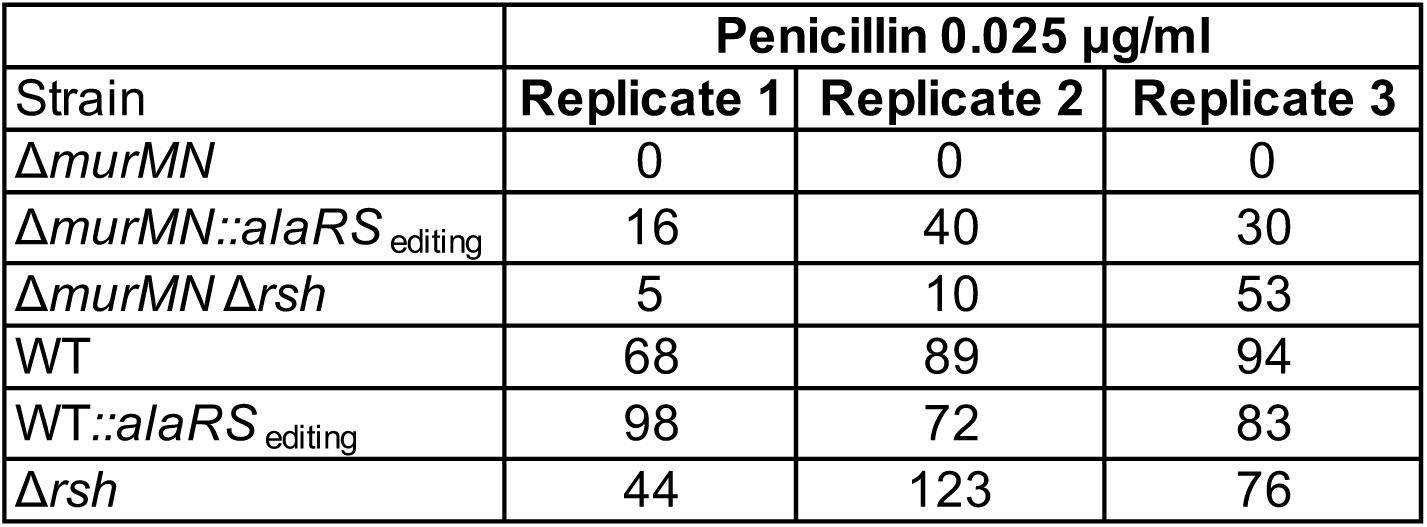
Percentage Viable CFUs of Different Strains after Growth in Penicillin G. Table represents percentage viability from three independent biological replicates.

|  | **Penicillin 0.025 µg/ml** |  |  |
| --- | --- | --- | --- |
| **Strain** | **Replicate 1** | **Replicate 2** | **Replicate 3** |
| *ΔmurMN* | 0 | 0 | 0 |
| *ΔmurMN::alaRS<sub>editing</sub>* | 16 | 40 | 30 |
| *ΔmurMN Δrsh* | 5 | 10 | 53 |
| WT | 68 | 89 | 94 |
| *WT::alaRS<sub>editing</sub>* | 98 | 72 | 83 |
| *Δrsh* | 44 | 123 | 76 |

### *murMN* dampen LytA-mediated lysis

The early onset of stationary phase is followed by increased lysis in the *ΔmurMN* strain when grown in mildly acidic conditions (**Fig. 1**). Having established that *murMN* regulate activation of the stringent response pathway, we searched for the molecule(s) responsible for the autolysis. Autolysis of pneumococcal cells is carried out by autolysins, peptidoglycan hydrolases containing choline-binding domains that enable their binding to phosphocholine residues present on the teichoic acids of the cell wall (38–40). Lysis requires attachment of choline-binding proteins to the cell surface, such that addition of exogenous choline inhibits this binding and the consequent autolysis (38, 41–43). Addition of two percent choline chloride to *ΔmurMN* grown in mildly acidic conditions abrogated lysis, suggesting lysis is triggered by choline binding protein(s) (**Fig. 5A**). Of the multiple choline binding proteins encoded in the pneumococcal genomes, LytA is the major autolysin. To investigate whether the increased lysis of *ΔmurMN* is induced by LytA, we tested growth of *ΔmurMN* with a deletion in *lytA* (*ΔmurMNΔlytA*). The *ΔmurMNΔlytA* strain resembled the wild-type strain in that it did not display lysis (**Fig. 5B**). Further, the lower maxOD observed in *ΔmurMN* strain was not rescued to wild type levels in the *ΔmurMNΔlytA* strain (**Fig. 5B**). This rescue of only the lysis phenotype in the *ΔmurMNΔlytA* strain suggests that in the absence of *murMN*, activation of the stringent response induces early onset of stationary phase and that LytA is responsible for the subsequent autolysis phenotype. The dramatically increased sensitivity of the *ΔmurMN* to LytA activity, relative to both the wild-type strain at mildly acidic and the *ΔmurMN* strain at normal conditions, suggest that *murMN* suppress the activation of LytA under mildly acidic conditions (**Fig. 5C**).

**Fig. 5.**
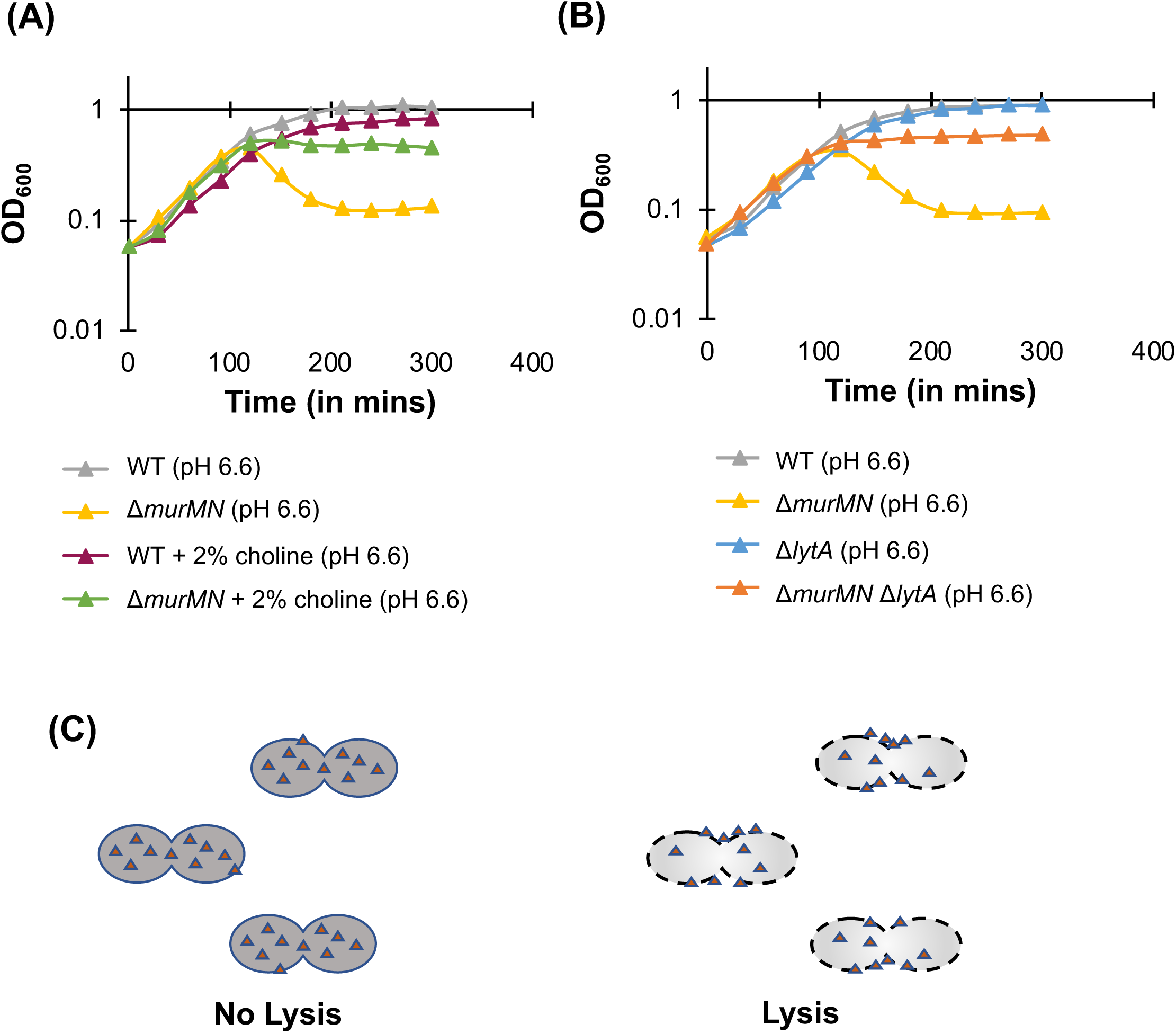
LytA-mediated autolysis is triggered in *ΔmurMN* cells. **(A)** Representative growth curve for wild-type and *ΔmurMN* strains grown in acidic conditions (▲), either in the absence or presence of 2% choline chloride. Growth curves were started at an OD_600_ of 0.05. **(B)** Representative growth curve for wild-type, *ΔmurMN, ΔlytA* and *ΔmurMNΔlytA* strains grown in acidic conditions (▲). Growth curves were started at an OD_600_ of 0.05. *(at least n=3)* **(C)** Schematic representing change in the distribution of LytA (triangles). When strains are growing in exponential phase, LytA is localized mainly intracellularly (*left*) but when they reach stationary phase, LytA is exported to the surface resulting in autolysis (*right*), based on findings from Mellroth *et al.* (66).

### MurMN influence macrophage phagocytosis by regulating LytA activity

Pneumococcal cells encounter numerous environmental stresses in an infection setting. Among these is acidic pH, encountered as a result of the inflammatory response mounted by the host against the invading bacteria (44, 45). Moreover, previous work suggests that LytA contributes to pneumococcal evasion from phagocytosis (46), and our data revealed that the *ΔmurMN* strain displays increased LytA mediated autolysis. Thus, we hypothesized that macrophages would be less efficient at uptaking the *ΔmurMN* strain relative to wild-type cells.

To test for phagocytosis, we moved to the encapsulated strain D39, the serotype 2 strain which is the ancestor of R6D. Macrophages were tested for their ability to phagocytose D39 and its isogenic mutants. In support of our hypothesis, the number of macrophages positive for pneumococcus was lower when infected with *ΔmurMN* strain relative to the wild-type strain (**Fig. 6, S1)**. Further, this phenotype was reversed for the *ΔmurMNΔlytA* strain, demonstrating the importance of LytA in this phenotype (**Fig. 6, S1)**. Finally, in agreement with previous work (46), the number of macrophages positive for pneumococcus was increased upon infection with a *ΔlytA* strain (**Fig. 6, S1)**.

**Fig. 6.**
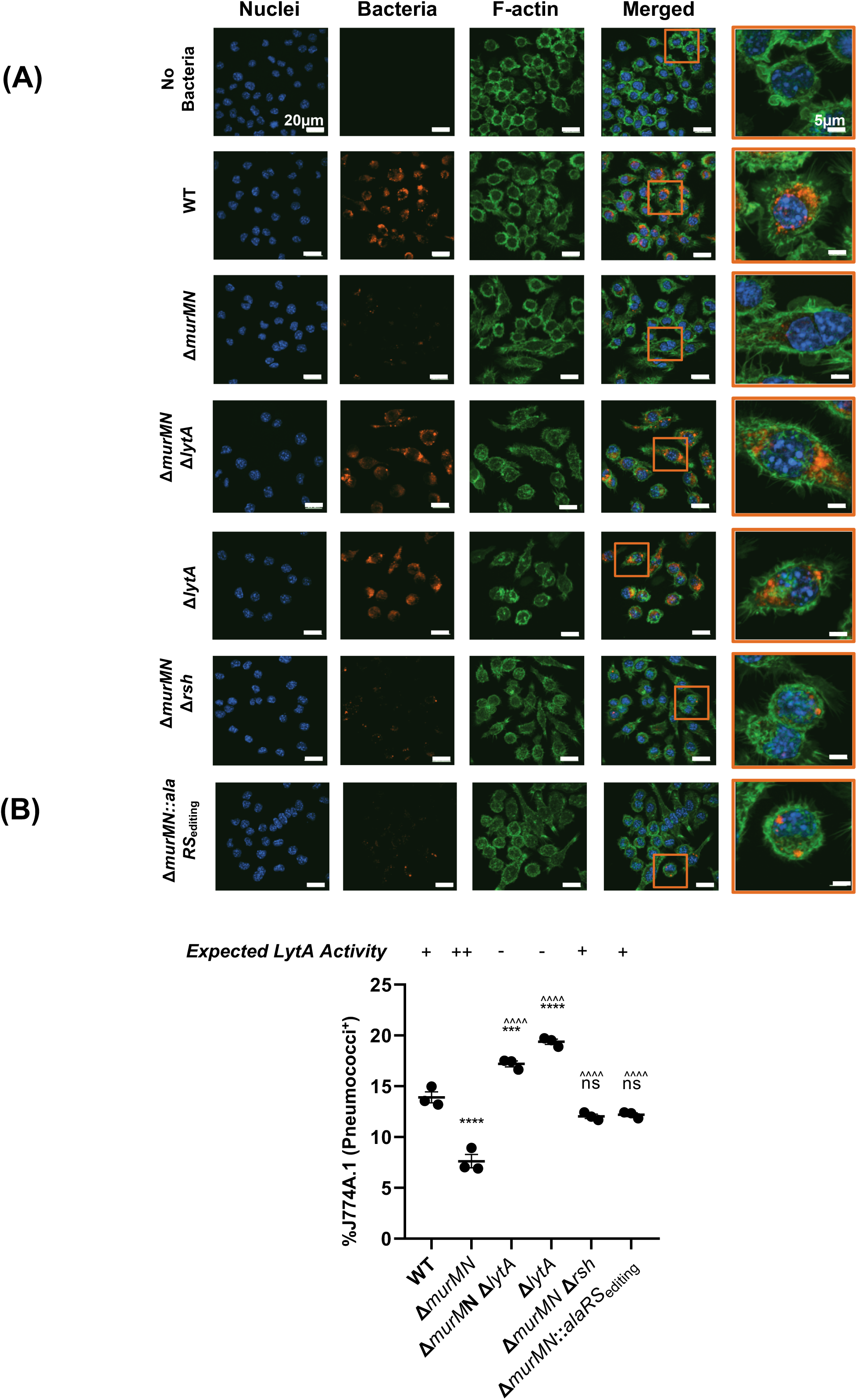
MurMN controls LytA-mediated evasion from phagocytosis. Internalization of pneumococcal cells with J774A.1 macrophages. **(A)** Confocal microscopy images depicting bacterial internalization after 60mins. *In blue*: macrophage nuclei, *red:* bacterial cells, *green*: actin. **(B)** Quantification of J774A.1 macrophages positive for pneumococci after 30mins, as separated by flow cytometry. ‘ns’ denotes non-significant comparisons, *** *p*<0.001, and \*\*\*\**p*<0.0001 relative to wild-type; ^^^^ *p*<0.0001 relative to *ΔmurMN*, by ANOVA followed by Tukey’s post-test. Graph also shows expected LytA activity for each strain, ‘+’ denotes basal levels, ‘++’ denotes increased levels and ‘-’ denotes absence.

Our data suggest that the activation of LytA in the *ΔmurMN* strain requires activation of the stringent response pathway. In support, the number of macrophages that internalize the *ΔmurMNΔrsh* strain was higher than those that internalize the *ΔmurMN* strain. Finally, our model predicts that *murMN* buffers the stringent response and LytA activation via its role in deacylating the misacylated tRNAs. Consistently, we observed a higher number of macrophages positive for the *ΔmurMN::alaRS*_editing_ strain relative to the *ΔmurMN* strain (**Fig. 6, S1)**. These data are consistent with a role for *murMN* in activation of the stringent response and consequent increase in LytA activity and capture the consequence of *murMN*-mediated stringent response suppression for phagocytosis.

The decreased percentage of macrophages positive for the *ΔmurMN* strain could be a consequence of either greater evasion of pneumococcal cells from phagocytosis or increased lysis of pneumococcal cells upon their internalization. If the acidic environment of macrophages promotes increased lysis of the *ΔmurMN* strain, there should be a rapid decrease in internalized bacteria over time. In contrast, we observe a more gradual decline in internalized *ΔmurMN* relative to the wild-type strain (**Fig. S2**). Thus, the *ΔmurMN* strain appears to evade phagocytosis, and, as previously reported, this process is LytA-dependent (46).

## DISCUSSION

Elucidating the mechanisms that bacterial cells employ to compartmentalize their functions or organize cell-wide responses to environmental stresses is fundamental to the field of microbiology. The tRNA-dependent-amino acid ligases involved in cell wall modification are optimally positioned to play a dual role in the cell: they build peptide crossbridges that serve as a major component of the bacterial cell wall, and they deacylate mischarged tRNAs, carrying out a vital role of translation quality control. Some bacterial species have evolved mechanisms to divert aminoacylated tRNA molecules away from translation and toward cell wall biosynthesis (12). For instance, *Staphylococcus aureus* encodes a set of isoacceptor tRNA species that display a greater propensity for peptidoglycan biosynthesis then translation, via diminished binding to the EF-Tu elongation factor utilized in translation (47). Similarly, *Mycobacterium tuberculosis* encodes dedicated tRNA synthetases that are spatially tethered to cell wall biosynthesis enzymes, and thus are exclusively used in the peptidoglycan (48). In contrast, in other bacterial species, the same pool of aminoacyl tRNAs serves both peptidoglycan modification and translation (49). Our data suggest that in this organization, cell wall tRNA-dependent-amino acid ligases can serve as regulators of intracellular stress. Our studies in the pneumococcus reveal that diminished editing and likely accumulation of mischarged tRNAs initiates the stringent response, premature entry into stationary phase, and subsequent activation of the murein hydrolase LytA. In this context, the cell wall modification protein MurM, by virtue of its role in deacylating mischarged tRNAs, dampens activation of the stringent response and calibrates the cellular reaction to stress.

An outstanding question in this model is how do mischarged tRNAs trigger the stringent response? Binding of aminoacyl tRNA to the ribosomal A-site provides the required amino acid for polypeptide elongation (16). In gram-negative bacteria, binding of deacylated tRNAs to the ribosome activates RelA and induces alarmone production (16). Activation of the stringent response is less understood in gram-positive bacteria, where the alarmone synthetase and hydrolase functions are fused in the bifunctional RSH enzyme (14, 50, 51). We propose that the accumulation of misacylated tRNAs, due to the absence of functional MurMN proteins, may also trigger the stringent response. Classic translation models suggest that the translation machinery recognizes the EF-Tu bound aminoacylated tRNA without displaying specificity for the amino acid, such that some amino acids from mischarged tRNAs can be incorporated into protein (52–55). However, other evidence suggest that the translation machinery recognizes both the amino acid and the tRNA: strong binding affinities of EF-Tu to tRNAs are compensated by weak binding affinities of EF-Tu to amino acids and vice-versa, leading to uniform binding affinities across tRNAs charged with their cognate amino acid (56). In this case, misacylated tRNAs can display binding affinities to the EF-Tu that are dramatically different than correctly charged tRNAs (57). Given the considerable difference in the inherent EF-Tu binding affinities of tRNA^Ala^ and tRNA^Ser^ (56, 58–60), it is expected that incorporation of Ser-tRNA^Ala^ will lead to an appreciable change in translation efficiency. Combined with our result that accumulation of mischarged tRNAs trigger the stringent response in pneumococcus, it seems plausible that RSH may be activated by binding of mischarged tRNAs to the ribosome.

Previous work has characterized the only RSH present in the pneumococcus, and established its important in alarmone production (37). The pneumococcal RSH contributes to copper uptake, provides increased fitness in cells subjected to mupirocin (an antibiotic that causes amino acid (Ile) starvation), and is a virulence determinant in the murine models of pneumococcal pneumoniae and sepsis (37). Further, studies of a RSH point mutation identified in a laboratory variant of a clinical strain, demonstrated that RSH promotes resistance to neutrophil killing and provides a fitness advantage in a murine model of colonization (61). Together, these studies strongly suggest that the ability of pneumococcus to trigger the stringent response is critical for virulence. We put forward the idea that MurMN play a role in fine-tuning entry into the stringent response, and as such may contribute to the outcome of pneumococcal infections. This could explain why most bacteria carry the *murMN* operon despite lacking a peptidoglycan rich in the branched muropeptides that are the product of the MurMN proteins.

Our work describing a novel role for *murMN* was performed using growth in a mildly acid condition. Bacterial cells commonly encounter acidic environment in the human host during infection due to inflammation response mounted by immune cells (45). There are a series of studies on molecular responses of pneumococcal cells subjected to acid stress (62–65). These studies describe the molecular regulation of acidic stress-induced lysis, a response mediated by LytA and regulated by phosphorylation of ComE in a competence stimulating peptide-independent manner (63–65). These studies differs from our work in design and scope. First, while we also focus on pneumococcal growth in acidic pH, these studies analyzed cells pre-conditioned in alkaline pH before exposure to acidic stress. Second, our work is centered on *murMN* and its role in the cellular response to acidic stress. The growth defect we describe is not observed in a *ΔcomE* strain and the growth of the *ΔmurMNΔcomE* strain resembles that of *ΔmurMN* strain (data not shown). Thus, the molecular responses described in this study do not appear to overlap with those previously described and highlights the importance of *murMN* in regulation of growth in mildly acidic conditions.

The *ΔmurMN* strain is prone to LytA-mediated lysis, and this effect is dramatic at a mildly acid pH. The dynamics of LytA during planktonic growth have been well characterized (66). During exponential growth, LytA is mainly intracellularly and inactive. During stationary phase, LytA exits the cells, where it promotes autolysis via its *N*-acetylmuramoyl-L-alanine amidase activity and in doing so breaks the amidase bond between the carbohydrate and the peptide components of the peptidoglycan backbone (67). Thus, the activation of LytA in the *ΔmurMN* strain can be explained by its early onset into stationary phase. In addition, previous work has demonstrated that LytA contributes to evasion from phagocytosis by inhibition of complement-mediated immunity (46). LytA inhibits binding of proteins in the classical complement pathway to the pneumococcal surface and enhances recruitment of suppressors of complement to impair the alternative complement pathway. Thus, we hypothesized that increased activity of LytA in the *ΔmurMN* strain would protect it from macrophage uptake. As predicted, the number of macrophages with internalized pneumococci was lower for the *ΔmurMN* relative to the wild-type strain. Moreover, this phenotype was reversed in a *ΔmurMNΔlytA* mutant. These data are consistent with our conclusion that *murMN* restrains LytA activity. Further, our experiments in macrophages combined with previous work on *Δrsh* strains (37, 61) suggest that pneumococcal fitness *in vivo* may depend on careful calibration of the stringent response, where activation and timing can expose or conceal pneumococcus to different facets of the immune response.

MurM is encoded in all pneumococcal strains as well as some related species, and there are multiple *murM* alleles (9, 68–70). All pneumococcal *murM* alleles have the ability to introduce both serine and alanine at the first position of the diamino acid peptide bridge (8). However, alleles vary regarding expression levels and the relative ratio of serine versus alanine they insert in the first position of the peptide bridge (8, 26). Thus, *murM* alleles may also vary in their ability to deacylate mischarged Ser-tRNA^Ala^ and control entry into the stringent response.

In nature, there is one free standing editing protein, AlaXp, and it deacylates Ser-tRNA^Ala^ and Gly-tRNA^Ala^. AlaXp is homologous to the editing domain of the AlaRS alanyl-tRNA synthetase protein, which possesses both synthetase and editing functions (33). The conserved widespread distribution of AlaXp across the tree of life contrasts with the limited phylogenetic distribution of other standalone aminoacyl-tRNA editing proteins and highlights the importance of the AlaXp function. Further, analysis of characterized AlaXp proteins reveals that they are conserved in their ability to deacylate Ser-tRNA^Ala^, but not necessarily Gly-tRNA^Ala^ molecules (33). These findings suggest that deacylating mischarged Ser-tRNA^Ala^ is of great importance to the cell, and that accumulation of this mischarged tRNA is toxic to the cell. Thus, it has been proposed that AlaXp is an evolutionary solution to the ubiquitous challenge of Ser-tRNA^Ala^ production. The overlap in function between MurM and AlaXp suggests that MurM may also represent an alternative evolutionary solution to the challenge of editing Ser-tRNA^Ala^ (34). Our genomic search reveals that many Firmicutes that produce peptide bridges, such as *Staphylococcus aureus*, *Streptococcus thermophilus*, *Streptococcus agalactiae*, *Streptococcus salivarius, Enterococcus faecalis* and *Lactobacillus viridescens*, do not appear to encode for an AlaXp protein. It is thus tempting to speculate that the composition of cell wall bridges, the vast majority of which utilize serine, alanine, or glycine, may represent an adaptation that reflects the function of cell wall tRNA synthetases in translation quality control.

In conclusion, this study suggests that MurM provides a molecular link between cell wall modification and translation quality control in gram-positive bacteria that produce short peptide bridges in their peptidoglycan layer. These bridges are structural components with a critical role in antibiotic resistance (6–8, 11). This study provides *in vivo* evidence to suggest that the propensity of MurM to deacylate mischarged-tRNA allows it to serve as a translation quality control checkpoint and dampen entry into the stringent response. Through these mechanisms, MurM regulates the activity of LytA and influences phagocytosis. These findings implicate MurM in the survival of bacteria as they encounter unpredictable and hostile conditions in the host.

The question of how the regulation of these fundamental biological processes by MurM influences the extent of pathogenesis still remains to be investigated. Our work provides functional insight into the role of peptidoglycan peptide bridge generating enzymes in modulating intracellular stress and entry into the stringent response.

## MATERIALS & METHODS

### Bacterial strains & growth conditions

This experimental work was performed on two strains of *Streptococcus pneumoniae* and their isogenic mutants. The majority of the work was performed on a penicillin-resistant derivative of R6 (R6D). This R6D isolate, also referred to as Hun663.tr4, was generated from a genetic cross where parental R6 strain was recombined with DNA isolated from Hungary19A-6 and recombinants were selected for penicillin resistance (71). The *murM* allele in this strain correspond to allele *murMB1*. R6D is a non-encapsulated strain, thus the macrophage experiments were performed with the highly related serotype 2 D39 strain (*murMA* allele) and isogenic mutants (GenBank CP000410) (72).

Colonies were grown from frozen stocks by streaking on TSA-II agar plates supplemented with 5% sheep blood (BD BBL, New Jersey, USA). Colonies were then picked and inoculated in fresh Columbia broth (Remel Microbiology Products, Thermo Fisher Scientific, USA) and incubated at 37°C and 5% CO_2_ without shaking. For acidic conditions, the pH of the Columbia broth was adjusted to 6.6 by the addition of 1M HCl. The basic recipe used for preparing chemically defined medium (CDM) was as previously described (73). MOPS-CDM was prepared by replacing potassium phosphate and sodium phosphate as previously described (37).

### Construction of mutants

The deletion mutant strains were constructed by using site-directed homologous recombination to replace the region of interest with antibiotic selection marker (Table S1). Briefly, the transformation constructs were generated by assembling the amplified flanking regions with the antibiotic resistance cassettes. Between 1-2kb of flanking regions upstream and downstream of the regions of interest were amplified from parental strains using Q5 2x Master Mix (New England Biolabs, USA). The antibiotic resistance gene *ermB* was amplified from *S. pneumoniae* SV35-T23, and the genes, *kan* and *aad9* were amplified from *S. pneumoniae* R6D*ΔbriC::briC* and R6D::spec^R^ (74), respectively. SV35-T23 has an *ermB*-containing mobile element, making it resistant to erythromycin (75). These PCR products were assembled together either by sticky-end ligation of restriction-cut PCR products or by Gibson Assembly using NEBuilder HiFi DNA Assembly Cloning Kit (New England Biolabs, USA).

The strains *ΔmurMN:murMN* and WT*::alaRS*_editing_ were generated by ligating the gene of interest at its 3’ end with an antibiotic resistance cassette. These were assembled with the amplified flanking regions either by sticky-end ligation of restriction-cut PCR products or by Gibson Assembly using NEBuilder HiFi DNA Assembly Cloning Kit. The *murMN* complement strain (*ΔmurMN:murMN*) was recombined at the native *murMN* locus. The WT*::alaRS*_editing_ was introduced downstream of *bga* (without modifying *bga*), a commonly employed site for transformation (74, 76). The *ΔmurMN::alaRS*_editing_ strain was generated by transformation of the corresponding *alaRS* construct into *ΔmurMN* strain. Primers used to generate the construct are listed in Table S2.

The D39 mutants (D39 *ΔmurMN,* D39 *ΔlytA*, D39*ΔmurMNΔlytA*, D39 *ΔmurMNΔrsh*, D39*ΔmurMN::alaRS*_editing_) were generated by transformation with the corresponding constructs amplified from R6D into strain D39.

### Bacterial transformations

To generate mutants, target strains (R6D or D39) were transformed in the following way. Strains were grown in acidic Columbia broth to an OD_600_ of 0.05, followed by the addition of 1μg of DNA along with 125μg/mL of CSP1 (sequence: EMRLSKFFRDFILQRKK; purchased from GenScript, NJ, USA). These treated cultures were incubated at at 37°C for 2 hours. Following the incubation, the cultures were plated on selective Columbia agar plates that contain the appropriate antibiotic: spectinomycin (100μg/ml), erythromycin (2μg/ml), or kanamycin (150μg/ml), and incubated at 37°C overnight. The following day, resistant colonies were selected and cultures in selective media, and colonies were confirmed using PCR. The bacterial strains generated in this study are listed in the Table S1.

### Growth curves

Strains of interest were streaked on TSA-II agar plates supplemented with 5% sheep blood (BD BBL, New Jersey, USA). These streaked cells were then inoculated into fresh Columbia broth (normal or acidic) and incubated at 37°C and 5% CO_2_ without shaking. Once the cultures reached an OD_600_ of 0.05, the growth of the cultures was followed every 30 minutes by recording their optical density at a wavelength of 600nm.

Strains with a *Δrsh* genetic background showed a much longer lag phase. To circumvent this, when comparing growth of cells with strains in *Δrsh* background, we adapted our protocol from a previous study measuring growth of these cells (37). The cells were grown in acidic Columbia broth to an OD_600_ of ~0.2. The cells were then collected by centrifugation at 3000 rpm for 10 minutes. These cells were then resuspended in acidic Columbia broth to an OD_600_ of approximately 0.1. The growth of these cultures was followed by every 30 minutes by recording their optical density.

All growth curves represented within a given figure panel were carried out in parallel and with the same batch of media.

### Cell wall preparation & enzymatic degradation

Pneumococcal cell walls were prepared following a previously published procedure (71). Briefly, after collecting cells by centrifugation, cells were suspended in ice-cold PBS and dropped into boiling sodium dodecyl sulfate to inactivate cell wall-modifying enzymes. The peptidoglycan was further purified using enzymatic degradation as previously described (6).

### Analysis of stem peptide composition

The stem peptides contained with the extracted and processed peptidoglycans were separated and analyzed by reverse-phase high-performance liquid chromatography (RP-HPLC) as has been previously described (6).

### Thin-layer chromatography (TLC)

*S. pneumoniae* strains were grown in acidic Columbia broth until an OD_600_ of 0.1 followed by an additional 1-hour incubation. This time-point was chosen so all cells were still in exponential phase, owing to the early lysis observed in *ΔmurMN* cells. The protocol for alarmone labelling and TLC was adapted from a previous study (37). Cells were harvested by centrifugation at 3300 *g* for 4 minutes. Pellets were washed and the cells were resuspended to the same density in MOPS-CDM. 200µL of the suspension was prewarmed at 37°C for 10 minutes. Following this, 25µL of the warmed suspension was mixed with 150µL of MOPS-CDM containing 150µCi mL^-1^ of H_3_[^32^P]O_4_ (2mCi mL^-1^ in Water; PerkinElmer, Waltham, MA) for 30 minutes. 25µL of the sample was then removed and mixed with an equal volume of frozen 13M formic acid, followed by repeated thawing and refreezing. After two such cycles, 5µL of the sample was removed for spotting for TLC. The samples were spotted on polyethyleneimine-cellulose (PEI) plastic-baked TLC plates (J.T. Baker) and chromatographed in 1.5M KH_2_PO_4_ (pH 3.4). TLC plates were then dried, and exposed to a phosphor screen (GE Healthcare, Chicago, IL) overnight. The plates were imaged using Typhoon FLA 9000 imager (GE Healthcare, Chicago, IL) and analyzed using ImageJ.

### Antibiotic fitness assay

Pneumococcal cells were grown to mid-exponential phase at an OD_600_ ~0.5 in fresh Columbia broth. Serial dilutions were performed and the cells were plated on either TSA-II blood agar plates or Columbia agar plates with 0.025 µg/ml of penicillin G, and incubated at 37°C and 5% CO_2_ overnight. Percent of viable cells was calculated as the ratio of viable CFUs on antibiotic plates relative to viable CFUs on blood agar plates.

### Phagocytosis assay

J774A.1 cells (#ATCC® TIB-67™, ATCC, Manassas, Virginia) were grown and maintained in Roswell Park Memorial Institute media (RPMI; Gibco, Gaithersburg, MD) supplemented with 10% fetal bovine serum (Thermo Fisher Scientific, Waltham, MA). J774A.1 cells were certified by IDEXX BioAnalytics (Columbia, MO) to be free of bacterial, fungal or mycoplasma contamination.

To test the phagocytosis of wild type and the different pneumococci mutants, 1×10^6^ J774A.1 cells were seeded per well in a tissue culture coated 6-well plate (Corning Inc, Corning, NY) and allowed to adhere overnight. Cells were infected in triplicate with 25μl of pneumococcal suspension containing ~3×10^7^ CFU resulting in a multiplicity of infection (MOI) of 10 and incubated at 37°C for 1 hour. Post incubation, cells were washed three times with PBS, and incubated for either 30, 60, 120 or 240 minutes in RPMI culture medium containing 10μg/ml penicillin and 200μg/ml gentamicin to kill extracellular pneumococci. Cells were then washed five times with PBS and phagocytosed pneumococci were harvested by treating the infected J774A.1 cells with 0.025% saponin for 15 minutes at 23 °C. Recovered bacteria were washed in PBS three times and assessed by flow cytometry. For flow cytometry, pneumococci cells were labelled with cell permeant SYTO13 green fluorescent DNA binding dye (Thermo Fisher Scientific, Waltham, MA) according to the manufacturer’s instruction prior to infecting J774A.1 cells. Infected J774A.1 cells were manually scrapped, washed and analyzed on Accuri C6 flow cytometer (BD Biosciences, San Jose, CA) connected to an Intellicyt HyperCyt autosampler (IntelliCyt Corp., Albuquerque, NM) using green channel (488 nm). Data were processed and interpreted using FlowJo software (FlowJo LLC, Ashland, Oregon).

### Confocal microscopy

J774A.1 cells were seeded on 12 mm collagen type-I coated glass coverslips (Electron Microscopy Services, Hatfield, PA) and infected with SYTO13 labeled wild type and the different pneumococci mutants with an MOI of 10 as described above. After 1 hour of infection with labelled pneumococci, J774A.1 cells were washed five times with PBS and treated with 10μg/ml penicillin and 200μg/ml gentamicin for 1 hour to kill extracellular bacteria. Cells were then washed three times with PBS and fixed with 3.33% freshly prepared paraformaldehyde (Electron Microscopy Services, Hatfield, PA) for 20 min at 23°C. Excess fixative was quenched by adding an equal volume of 1% (w/v) BSA in PBS for 5 min followed by three washes with PBS. To visualize F-actin and nuclei, cells were stained with Alexa Fluor™ 647-Phallodin (Thermo Fisher Scientific, Waltham, MA) (5:200 in PBS) and Hoechst 33342 (Thermo Fisher Scientific, Waltham, MA) (1:1000 in PBS), respectively. Imaging was performed using a Carl Zeiss LSM 880 confocal microscope with fixed settings and the images were analyzed using ZEN Black software (Carl Zeiss Microscopy, Thornwood, NY).

### Statistical tests

The statistical differences were calculated using ANOVA. Comparison among different groups was performed using Tukey’s post-test. *p* values of less than 0.05 were considered to be statistically significant.

### Analysis of the *S. pneumoniae* genomes for AlaXp

A Blastp search was implemented in NCBI. As query we use the AlaXp sequences from Bacillus sp (GenBank WP_000206094.1). Using default parameters, we screened the following NCBI genomes: (1) *Staphylococcus aureus (taxid:1280), Streptococcus thermophilus (taxid:1308), Streptococcus agalactiae (taxid:1311)*, *Streptococcus salivarius (taxid:1304), Enterococcus faecalis (taxid:1351)*, and *Lactobacillus viridescens (taxid:1629)*. All top hits had p-value below 1e-15 and approximately 30% percent identity to the query sequence, except for *Weissella viridescens* and some Enterococci strains that displayed p-value ~1e-9 and percent identity around 25%. Hits were analyzed to establish the starting residue on the query. In all these genomes this number ranged from 489-499, consistent with AlaRS. None of these genomes revealed a hit that matched the AlaXp query in size as well as sequence (although a few partial sequences were identified that appear to correspond to fractions of the AlaRS).

## ACKNOWLEDGEMENTS

We thank Prof. Alexander Tomasz for introducing us to the study of MurMN through stimulating discussions and challenging questions. We also thank him for providing the R6D strain used in this study. We thank Rory Eutsey and Karina Mueller-Brown for their assistance with laboratory experiments, Manning Huang for help with data visualization, Dr. Aaron Mitchell for productive advice throughout the development of this project, and Dr. John Woolford for a careful reading of this manuscript.

