## Supplementary Figures for "A Molecular Link between Cell Wall Modification and Stringent Response in a Gram-positive Bacteria"

### Figure S1

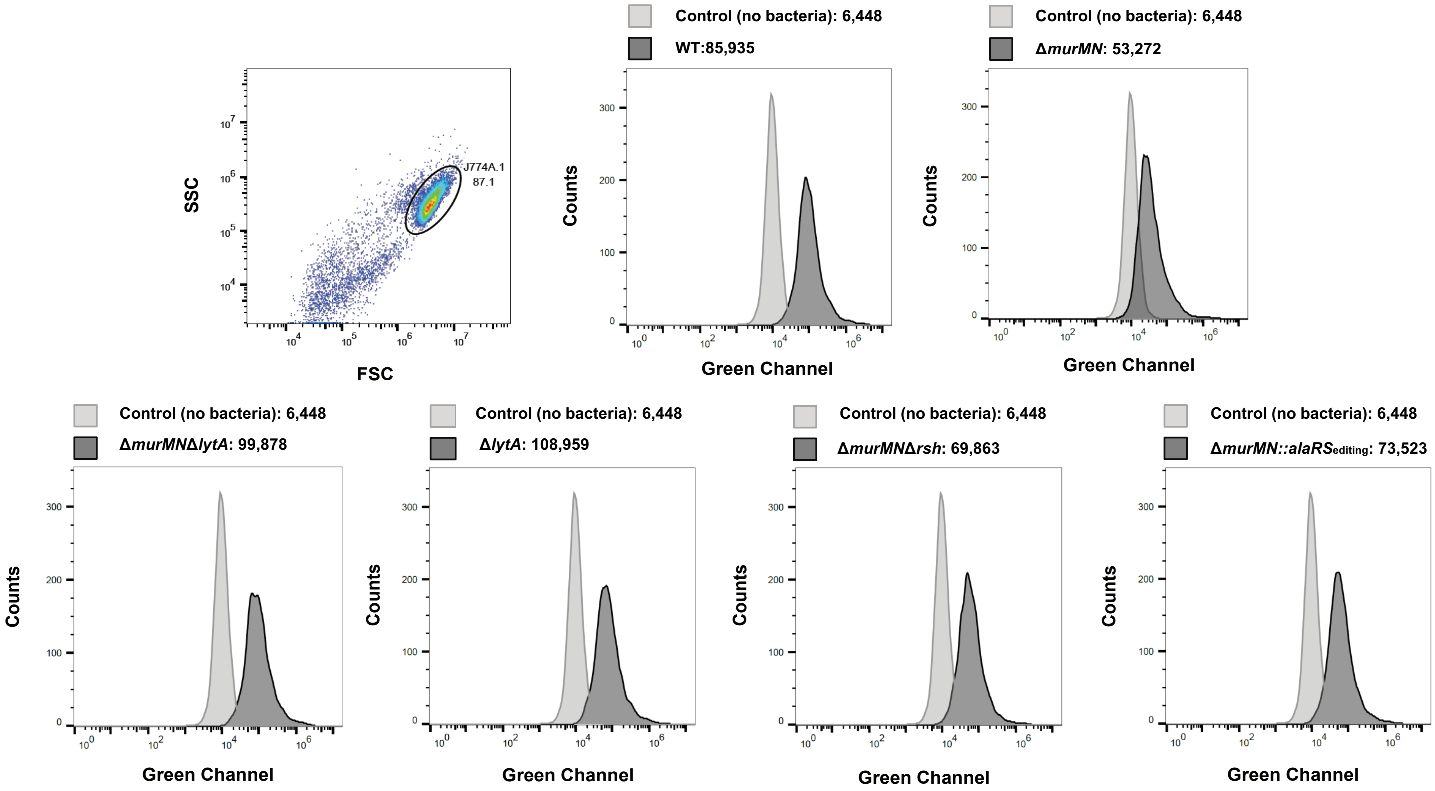


**Fig. S1. MurMN regulates pneumococcal internalization within J774A.1 macrophages.** Representative histograms depicting internalization of pneumococcal wild-type**,** *ΔmurMN, ΔmurMNΔlytA, ΔlytA, ΔmurMNΔrsh,* and *ΔmurMN::alaRS*$_{\text{editing}}$ strains with J774A.1 macrophages after 30mins using flow-cytometry. Each histogram depicts two fluorescence peaks – macrophages with no bacteria & macrophages infected with one pneumococcal strain. The mean fluorescence intensity values is also depicted. The first panel shows the forward versus side scatter used for gating the population of interest (encircled).

### Figure S2

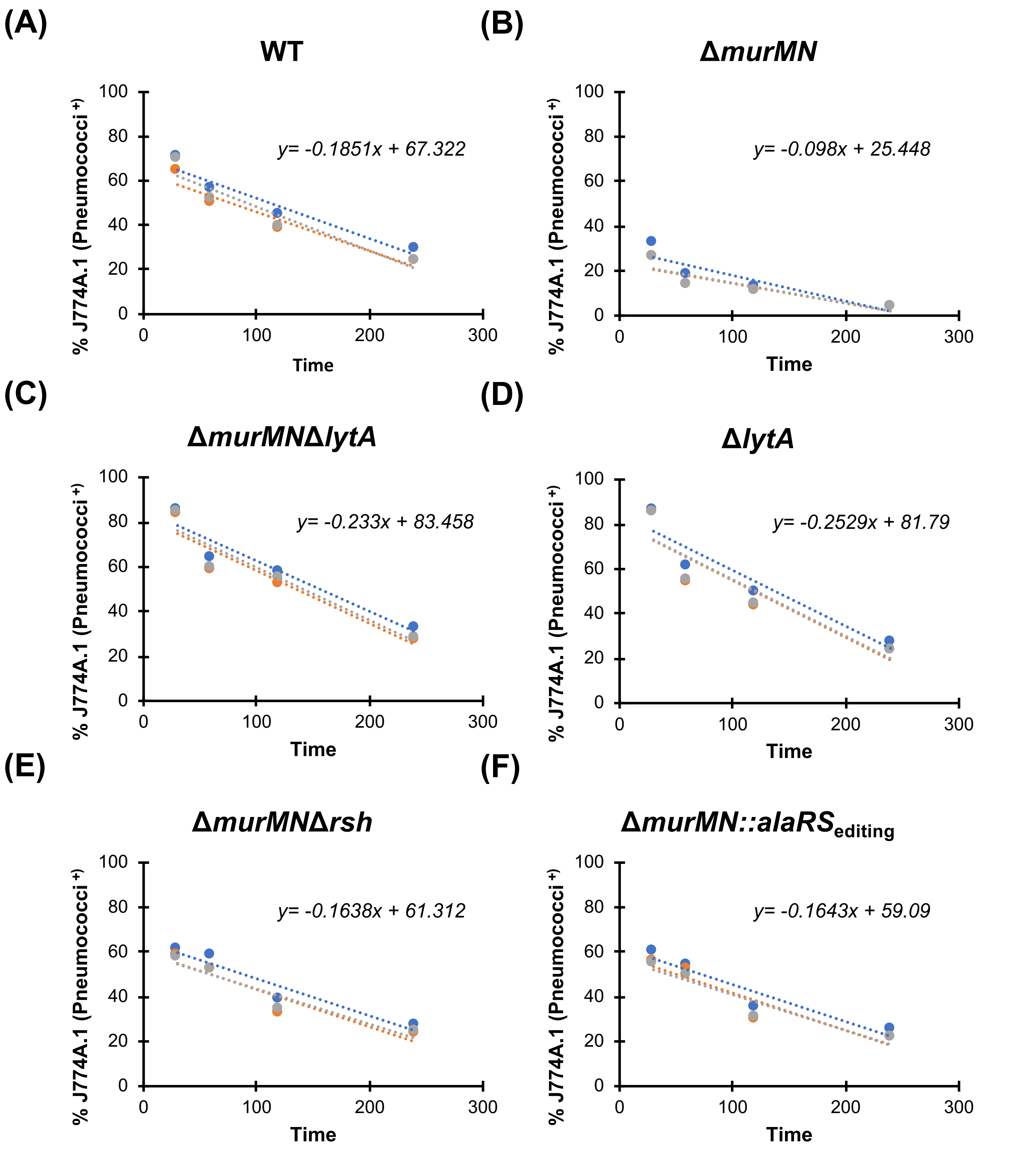


**Fig. S2. Bacterial internalization in macrophages over time.** Internalization of pneumococcal **(A)** wild-type **(B)** Δ*murMN* **(C)** Δ*murMN*Δ*lytA* **(D)** Δ*lytA* **(E)** Δ*murMN*Δ*rsh* **(F)** Δ*murMN::alaRS*$_{\text{editing}}$ strains with J774A.1 macrophages for 30, 60, 120, 240mins. Each data point represents percent macrophages positive for pneumococcal cells at the indicated time. The dotted line represents the linear fit trendline indicating decline in percent positive macrophages. Each panel consists of three biological replicates depicted in different colors (blue, orange, grey). The linear fit trendline of the average of the three replicates is defined by the written equation of the type *y=mx+c*, where 'm' is the slope, and 'c' is the intercept.
